# Three-dimensional morphological analysis revealed the cell patterning bases for the sexual dimorphism development in the liverwort *Marchantia polymorpha*

**DOI:** 10.1101/2023.01.24.525312

**Authors:** Yihui Cui, Tetsuya Hisanaga, Tomoaki Kajiwara, Shohei Yamaoka, Takayuki Kohchi, Tatsuaki Goh, Keiji Nakajima

**Affiliations:** Graduate School of Science and Technology, Nara Institute of Science and Technology, 8916-5 Takayama, Ikoma, Nara 630-0192, Japan; Graduate School of Biostudies, Kyoto University, Kitashirakawa Oiwake-cho, Sakyo-ku, Kyoto 606-8502, Japan

**Keywords:** Cell patterning, Gametophyte, Germline, Sexual dimorphism, Sexual reproduction, *Marchantia polymorpha*

## Abstract

In land plants, sexual dimorphism can develop in both diploid sporophytes and haploid gametophytes. While developmental processes of sexual dimorphism have been extensively studied in the sporophytic reproductive organs of model flowering plants such as stamens and carpels of *Arabidopsis thaliana*, those occurring in gametophyte generation are less well characterized due to the lack of amenable model systems. We here performed three-dimensional morphological analyses of gametophytic sexual branch differentiation in the liverwort *Marchantia polymorpha*, using high-depth confocal imaging and a computational cell segmentation technique. Our analysis revealed that specification of germline precursors initiates in a very early stage of sexual branch development where incipient branch primordia are barely recognizable in the apical notch region. Moreover, spatial distribution patterns of germline precursors differ between males and females from the initial stage of primordium development in a manner dependent on the master sexual differentiation regulator Mp*FGMYB*. In later stages, distribution patterns of germline precursors predict the sex-specific gametangia arrangement and receptacle morphologies seen in mature sexual branches. Taken together, our data suggests a tightly coupled progression of germline segregation and sexual dimorphism development in *M. polymorpha*.

## Introduction

Sexual dimorphism is commonly observed in multicellular organisms that propagate by sexual reproduction (Barrett and Hough 2013; McPherson and Chenoweth 2012). In oogamous animals, sexual dimorphism occurs in size and motility of haploid gametes, with female and male producing large immotile eggs and small motile sperm, respectively. In addition to the gamete morphology, sexual dimorphism also develops in diploid reproductive organs, and they play important roles in supporting gamete production, fertilization, and embryo development. As in animals, multicellular plants also undergo sexual differentiation in both reproductive organs and gametes. However, as land plants develop multicellular bodies both in haploid and diploid generations called gametophyte and sporophyte, respectively, plants exhibit additional sexual dimorphism in the gametophyte generation (Coelho et al. 2018; Schmidt et al. 2015). In flowering plants where sporophyte generation is predominant in the life cycle, conspicuous sexual dimorphism develops in the floral organs such as stamens and carpels, whereas sexual differentiation in the haploid gametophytes is inconspicuous, making pollens and embryo sacs that are composed of only a few cells and develop inside the sporophytic sexual organs (Schmidt et al. 2015). Thus, to study the process and mechanisms of gametophytic sexual differentiation in plants, a good model system with a gametophyte-dominant life cycle is required.

The recently reviving model bryophyte *Marchantia polymorpha* has been widely used as a unique system to study gametophytic sexual reproduction and their evolution in the land plant lineage (Hisanaga et al. 2019b). The life cycle of *M. polymorpha* is dominated by the haploid gametophytic generation. The main vegetative body consists of a leaf-like organ called thallus, which bifurcates at the apical notch region located between the bases of the two lobes (Shimamura 2016). *M. polymorpha* is dioicous, with its sex of its haploid gametophytes determined by the presence of either V or U chromosomes, yet no distinct sexual morphologies develop during vegetative growth (Iwasaki et al. 2021; Kohchi et al. 2021; Shimamura 2016; Yamato et al. 2007). In experimental cultures, reproductive growth of *M. polymorpha* is induced by supplemental irradiation of far-red (FR) light (Inoue et al. 2019). After phase transition, a sexual branch (gametangiophore) is formed from one of the two apical notch regions in a bifurcating thallus. Gametangiophores of *M. polymorpha* exhibit clear sexual dimorphism in their receptacle (Figure 1A and 1B). In the male gametangiophore (antheridiophore), receptacles exhibit a disc-like morphology with typically eight lobes per receptacle, whereas receptacles of female gametangiophores (archegoniophores) bear 9-11 finger-like rays (Shimamura 2016). In addition to the receptacle morphology, mature antheridiophores and archegoniophores exhibit clear difference in the spatial arrangement of gametangia in the receptacles (Figure 1C-1F). Male gametangia (antheridia) are embedded in the upper surface of the receptacles, whereas female gametangia (archegonia) are radially aligned beneath the receptacles. Moreover, developing gametangia are arranged in opposite orientations along the receptacle radius between males and females; more mature antheridia are located toward the center of the male receptacles, whereas more mature archegonia are located toward the periphery of female receptacles (Figure 1E and 1F) (Shimamura 2016).

**Figure 1.**
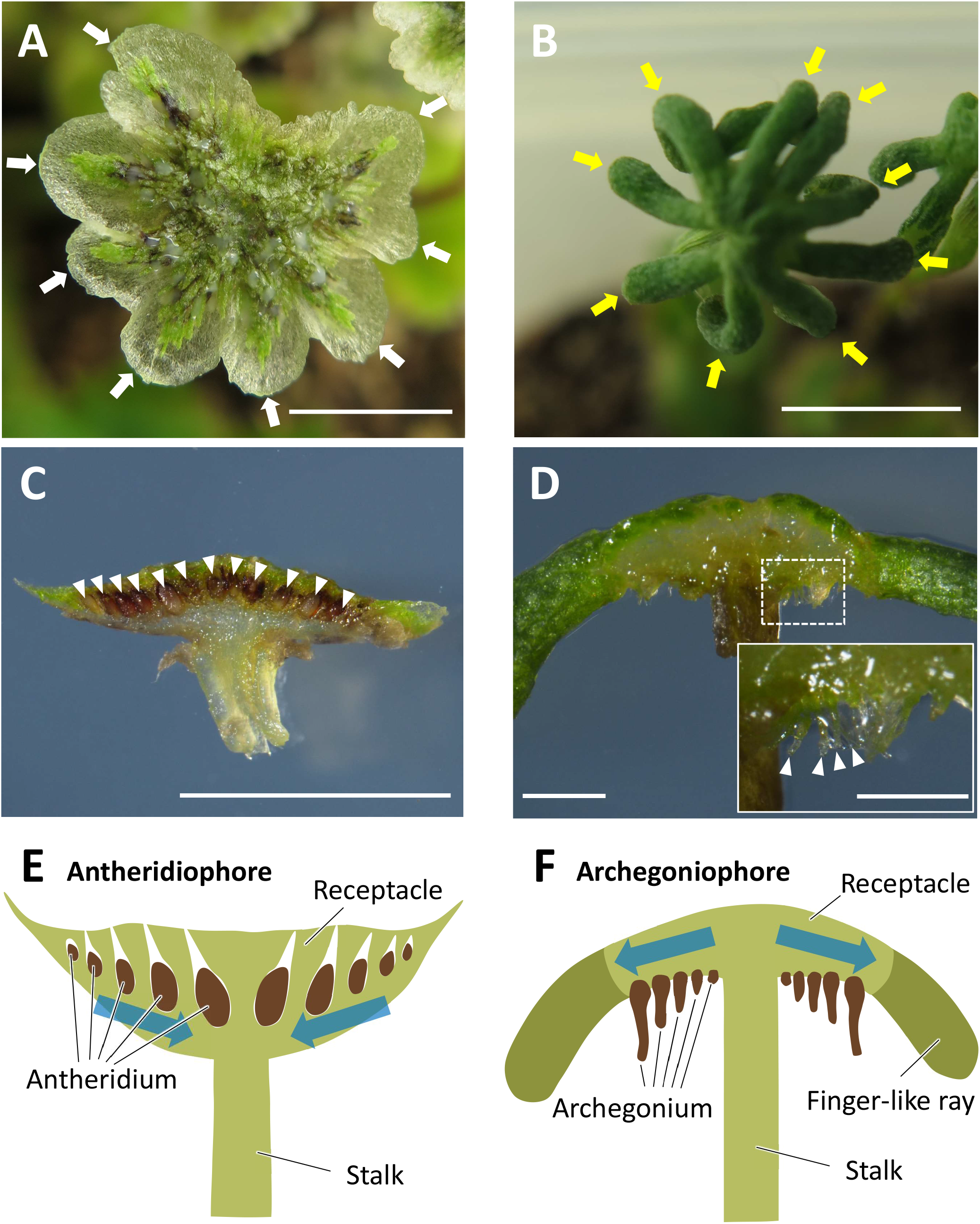
Morphologies of mature antheridiophore and archegoniophore of *M. polymorpha*. **(A, B)** Top view of antheridiophore (A) and archegoniophore (B). White and yellow arrows indicate lobes (A) and finger-like rays, respectively. **(C, D)** Cross sections of antheridiophore (C) and archegoniophore (D). White arrowheads indicate antheridia (C) and archegonia (D), respectively. Inset in (D) is a magnification of the boxed area. **(E, F)** Schematic drawings of an antheridiophore (E) and an archegoniophore (F). Note that the size of antheridia and archegonia are not to the real scale. Blue arrows indicate the direction of progression of gametangium maturation. Scale bar, 5 mm (A, B); 1 mm (C, D); 500 µm, inset in (D).

Previous studies have identified key regulators of sexual reproduction in *M. polymorpha*. Among them, Mp*BONOBO* (Mp*BNB*) encoding a member of the VIIIa subfamily of basic helix-loop-helix transcription factors promotes sexual branch formation and germ cell differentiation (Yamaoka et al. 2018). Mutants constitutively expressing Mp*BNB* form sexual branches independently of FR irradiation, whereas Mp*bnb* knock-out mutants are unable to form sexual branches even under prolonged FR irradiation. Expression of Mp*BNB* is transiently observed in gametangium initials that later give rise to gametangia in both males and females. *A. thaliana* mutants lacking two Mp*BNB* homologs are defective in generative cell specification in pollens, and this defect was rescued by Mp*BNB* expressed from the *A. thaliana BNB2* promoter, indicating evolutionarily conserved functions of *BNB* in germline differentiation (Yamaoka et al. 2018). Autosomal *FEMALE GAMETOPHYTE MYB* (Mp*FGMYB*) encoding an R2R3 MYB-type transcription factor is a key regulator of female sexual differentiation in *M. polymorpha* (Hisanaga et al. 2019a). Genetically female Mp*fgmyb* knock-out mutants exhibit a nearly complete female-to-male sexual conversion phenotype; they produce male sexual branches and sperm-containing antheridia, though sperm motility is lost presumably due to the lack of gene(s) in the V chromosome. Expression of Mp*FGMYB* is suppressed in males by the *cis*-acting anti-sense long non-coding RNA gene named *SUPPRESSOR OF FEMINIZATION (SUF)* in the Mp*FGMYB* locus. Loss-of-function *suf* mutant males exhibit female morphologies, though egg cells do not mature in their feminized gametangia. Expression of *SUF* is in turn silenced in females by U chromosomal Mp*BPCU* encoding a member of the BASIC PENTACYSTEINE transcription factor family (Iwasaki et al. 2021). Mp*BPCU* and its male gametolog Mp*BPCV* are also required for phase transition. Mp*FGMYB* is phylogenetically related to *A. thaliana MYB64, MYB98*, and *MYB119* known to regulate embryo sac and synergid cell differentiation (Kasahara et al. 2005; Rabiger and Drews 2013). Thus at least some members of the FGMYB clade appear to have roles in female gametophyte differentiation along the land plant lineage (Hisanaga et al. 2019a).

While *M. polymorpha* is a widely used bryophyte model and proven useful to decipher genetic mechanisms and evolution of sexual reproduction in land plants (Hisanaga et al. 2019b), detailed developmental processes of its sexual dimorphism, especially those leading to distinct sexual branch morphologies, have not been explicitly described at the cellular resolution, though cell division sequences to produce the female and male gametangia, i.e. archegonia and antheridia, have been documented for more than a century (Durand, 1908). In this study, we performed detailed three-dimensional (3D) morphological analysis to reveal cell-level developmental processes underlying the sexual dimorphism development in *M. polymorpha*. Using Mp*BNB* as a marker for germline precursors (Yamaoka et al. 2018), we visualized the spatial arrangement of germline segregation and their contribution to the sex-specific receptacle morphologies. Our study revealed that specification of germline precursors starts as early as in the stage where the sexual branch formation is barely detectable at the apical notch region. Moreover, spatiotemporal distribution patterns of germline precursors differ between male and female before the sex-specific organ morphologies become evident, and are later translated into the sex-specific gametangium arrangement and sexual branch morphologies, suggesting a link between germline positioning and sexual organ morphogenesis in *M. polymorpha*.

## Results

### Germline specification and gametangiophore development initiate simultaneously in *M. polymorpha*

To elucidate the cell-level patterning processes underlying the sexual dimorphism development in *M. polymorpha*, we performed 3D cell segmentation analyses of early-stage gametangiophore primordia. Briefly, segments containing gametangiophore primordia were excised from the apical notch region of thalli, subjected to tissue clearing by iTOMEI (Sakamoto et al. 2022), and stained with the fluorescence dye Renaissance 2200 to visualize cell walls. 3D cell wall patterns were reconstructed from Z-stack confocal images and used for cell segmentation analysis by the MorphoGraphX program (Figure 2A-2D) (Strauss et al. 2022). The primordia analyzed in this study were less than 500 µm in diameter and yet without elongated stalks. Thus, our analysis further dissected the previously defined stage 1 (< 2 mm in receptalce diameter) (Higo et al. 2016) into substages 1a through 1e (1a, <100 µm; 1b, 100-200 µm; 1c, 200-300 µm; 1d, 300-400 µm; 1e, >400 µm). We utilized the Mp*BNB*-*Citrine* knock-in lines to detect initiation of gametangia primordia (Yamaoka et al. 2018). Consistent with the previous report (Yamaoka et al. 2018), expression of MpBNB-Citrine was confined to the gametangium initials and their daughter cells (Figure 2G). Because these cells are segregated to sperm-forming antheridia and egg-forming archegonia, we here call the Mp*BNB*-Citrine-expressing cells germline precursors (colored pink in segmented images) (Lanfear 2018; Schmidt et al. 2015). Cells derived from the germline precursors could be easily identified by their characteristic cellular patters forming gametangium primordia (colored purple in segmented images). Thus, combination of the Mp*BNB* marker and cellular patterns allowed us to identify cell lineages leading to gametangium formation (Figure 2E-2G).

**Figure 2.**
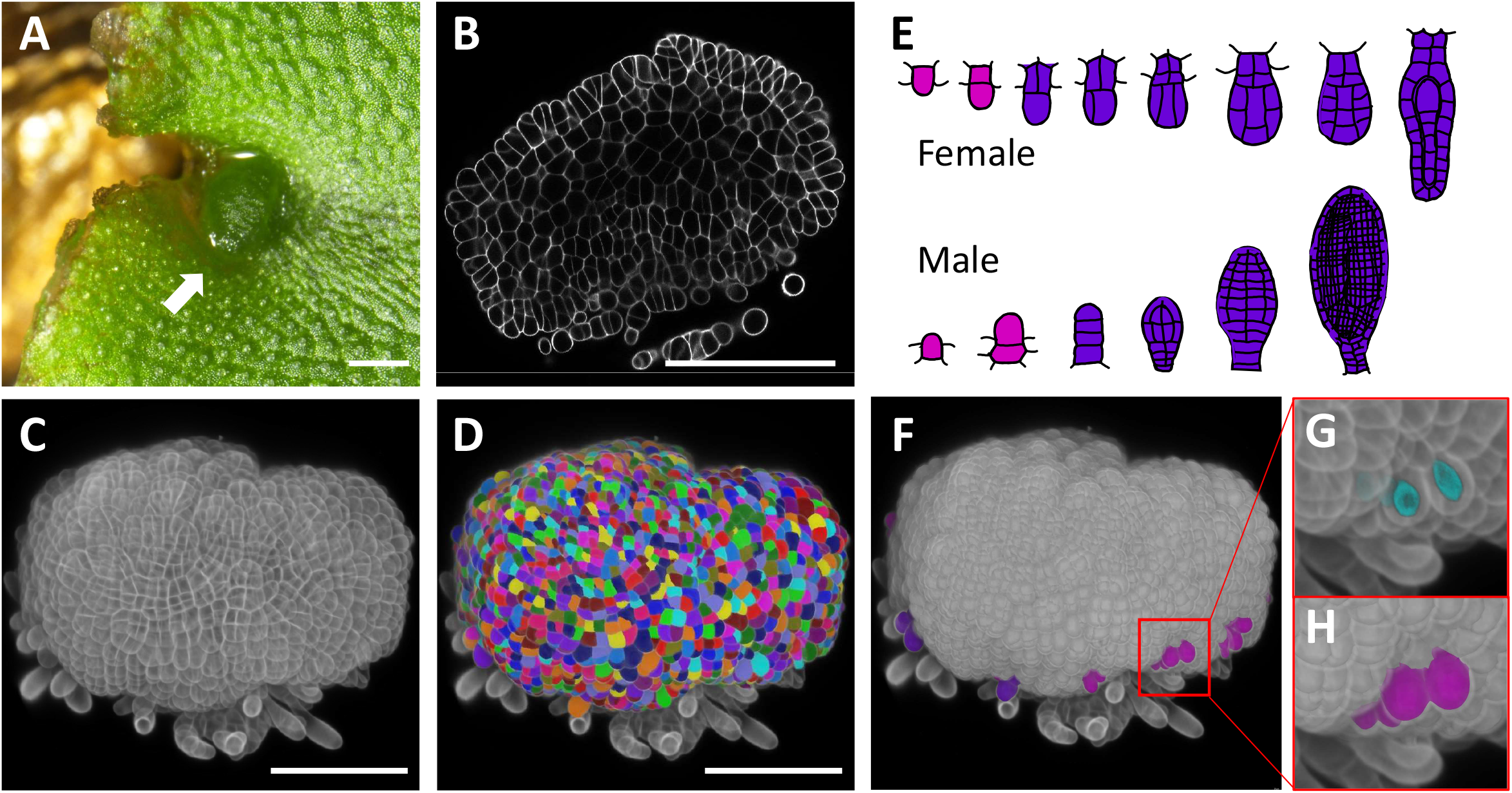
Outline of the 3D morphological analysis of gametangiophore primordia performed in this study. **(A)** A representative picture of an archegoniophore primordium developing in the apical notch region (arrow). **(B)** A representative confocal section showing the R2200-stained cell wall pattern of an archegoniophore primordium. **(C)** Color scheme of germline cells leading to the archegonium (top) and the antheridium (bottom). Germline lineage was identified based on the expression of the Mp*BNB-Citrine* reporter in early stages (pink) and on their characteristic morphologies in later stages (purple). **(D-H)** Outline of the 3D segmentation. Z-stack images were used to reconstruct 3D images (D), which were then used for the 3D-cell segmentation by MorphoGraph X (E). Germline precursors defined by the scheme shown in (C) were annotated on the segmented image (F). (G) and (H) are magnified images of the boxed region in (F). Original fluorescent image of Mp*BNB-Citrine* (G, cyan) and color-annotated image (H, pink) are shown. Scale bar, 500 µm (A); 100 µm (B-F); and 5 µm (G, H).

In males, visible antheridiophore primordia emerged as a slightly convex protrusion in the apical notch region typically two days after the onset of the FR irradiation, and a few MpBNB-expressing germline precursors were scattered over the primordium surface (Figure 3A and 4, stage 1a; Supplementary video 1). In later stages, more germline precursors emerged on the expanded primordia (Figure 4, stage 1b). In females, although the MpBNB-expressing germline precursors emerged in the incipient receptacle primordia as in males and in a slightly later timing (3 days after the onset of the FR irradiation), their number remained few in early stages (Figure 3B and 5, stages 1a and 1b; Supplementary video 2). Together, our observation confirmed the specific expression of MpBNB-Citrine in germline precursors (Yamaoka et al. 2018), and further revealed previously undescribed difference in the distribution patterns of germline precursors between male and female.

**Figure 3.**
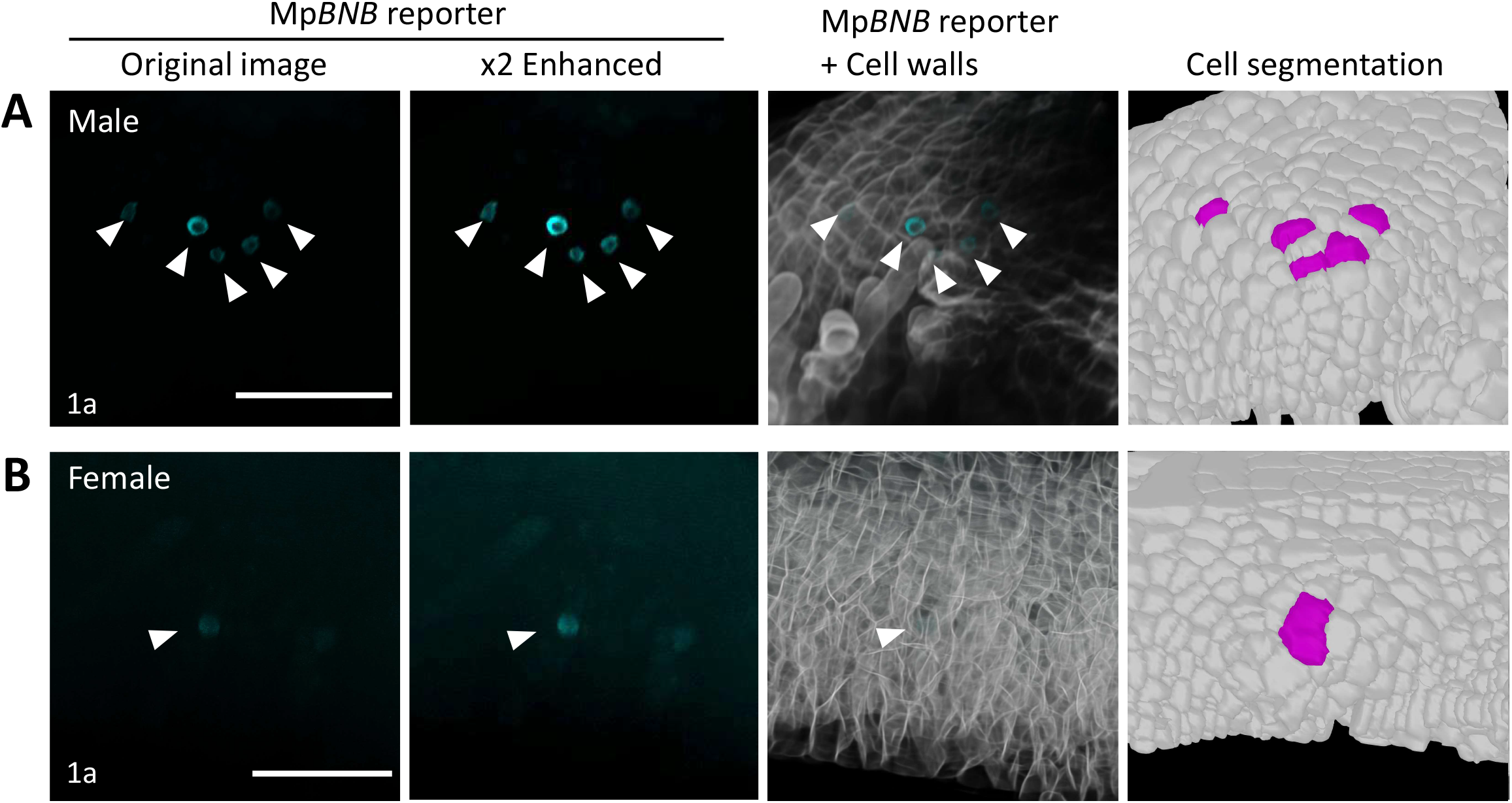
Specification of MpBNB-expressing germline precursors in the apical notch region after the induction of reproductive growth by FR. In both males (A) and females (B), MpBNB-expressing germline precursors emerge on the convex protrusion of incipient gametangiophore primordia (arrowheads). In males, multiple germline precursors emerge (A), whereas in females, only one germline precursor emerges. For better visualization, 2x enhanced images (equal enhancement for all RGB channels in the entire image area) of the original ones (left) are shown in the second column. Scale bar, 50 µm.

**Figure 4.**
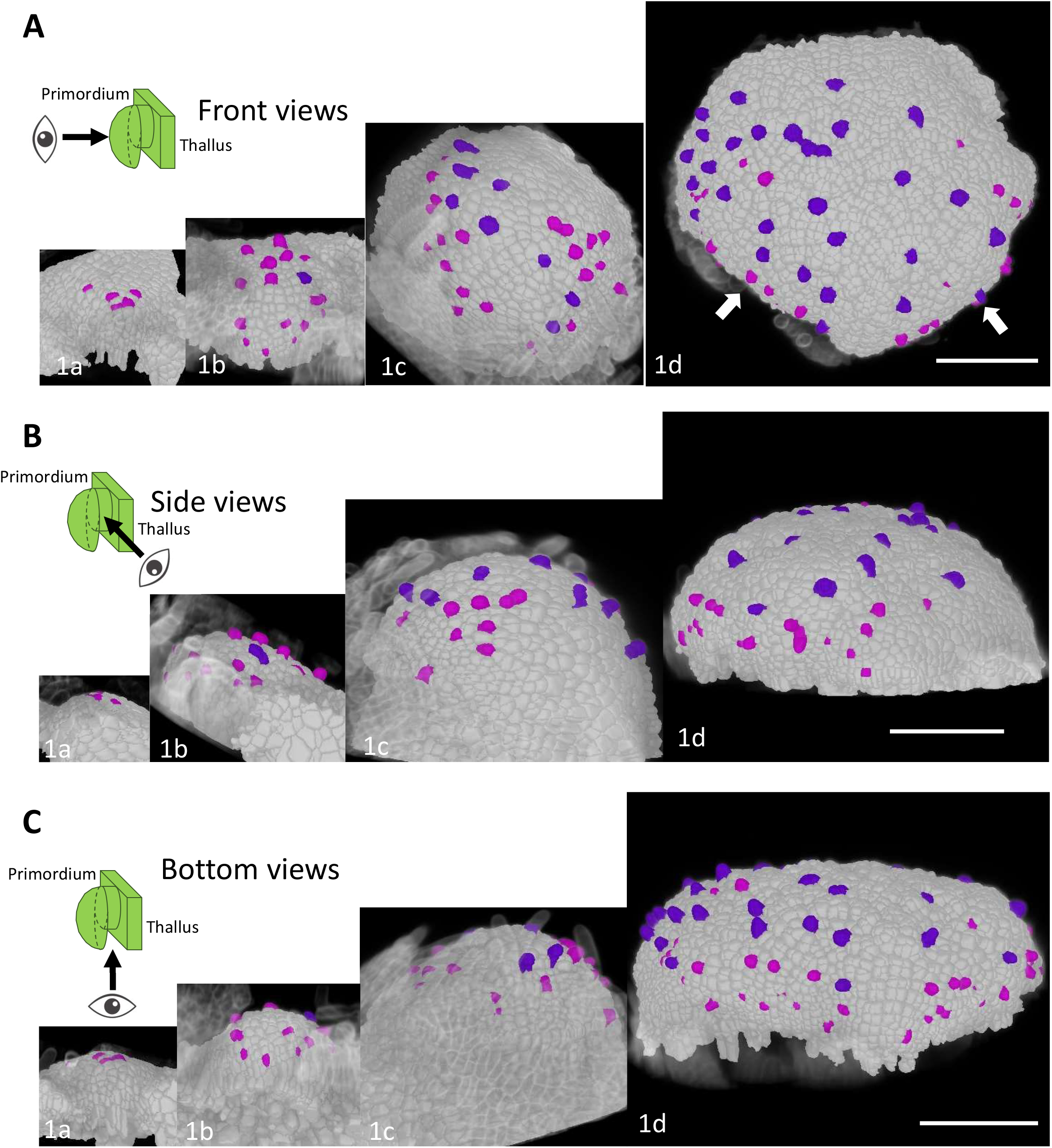
Morphogenesis and germline positioning during the antheridiophore development. Front views (A), side views (B) and bottom views (C) are shown for the primordia of different stages (1a-1d). Germline cells of early and late stages are colored pink and purple, respectively, according to Fig. 2E. White arrows in (A) indicate incipient lobes. Primordia of stages 1a, 1b, 1c, and 1d were taken 2, 3, 4, and 5 days after the onset of FR irradiation, respectively. Scale bar, 100 µm.

### Morphology of gametangiophore primordia and germline positioning differentiate between males and females early in the primordium development

While both male and female gametangiophore primordia initially had a similar dome-like morphology, they differentiate as the primordia grew larger in later stages. The overall morphology of male primordia became flattened (Figure 4, stage 1d; Supplementary video 3), whereas female primordia remained dome-shaped (Figure 5, stage 1d; Supplementary video 4). Difference in the number and distribution patterns of germline precursors also became more evident. In males, germline precursors and their progenies were broadly distributed over the upper surface of antheridiophore primordia (Figure 4 stages 1c and 1d; Supplementary video 3). This scattered distribution pattern was maintained as the primordia expand, while new antheridial initials continuously emerged at the peripheral region. Lobe-like protrusions emerged around disc-shaped primordium periphery (Figure 4, stage 1d). In later stages, germline precursors and their progenies were distributed around the incipient peripheral lobes of receptacle primordia, whereas more mature antheridial primordia were located closer to the receptacle center (Figure 4, stage 1d; Supplementary video 3). In females, the number of germline precursors remained fewer than those in males (Figure 6G). In contrast to males that produce lobe-like protrusion at the receptacle periphery (Figure 7A; Supplementary video 7), female primordia produced indentations on the bottom side of the receptacles (Figure 5, stage 1d; Supplementary video 4), which later became the indentations separating the finger-like rays (Figure 7B and 7C; Supplementary video 8).

**Figure 5.**
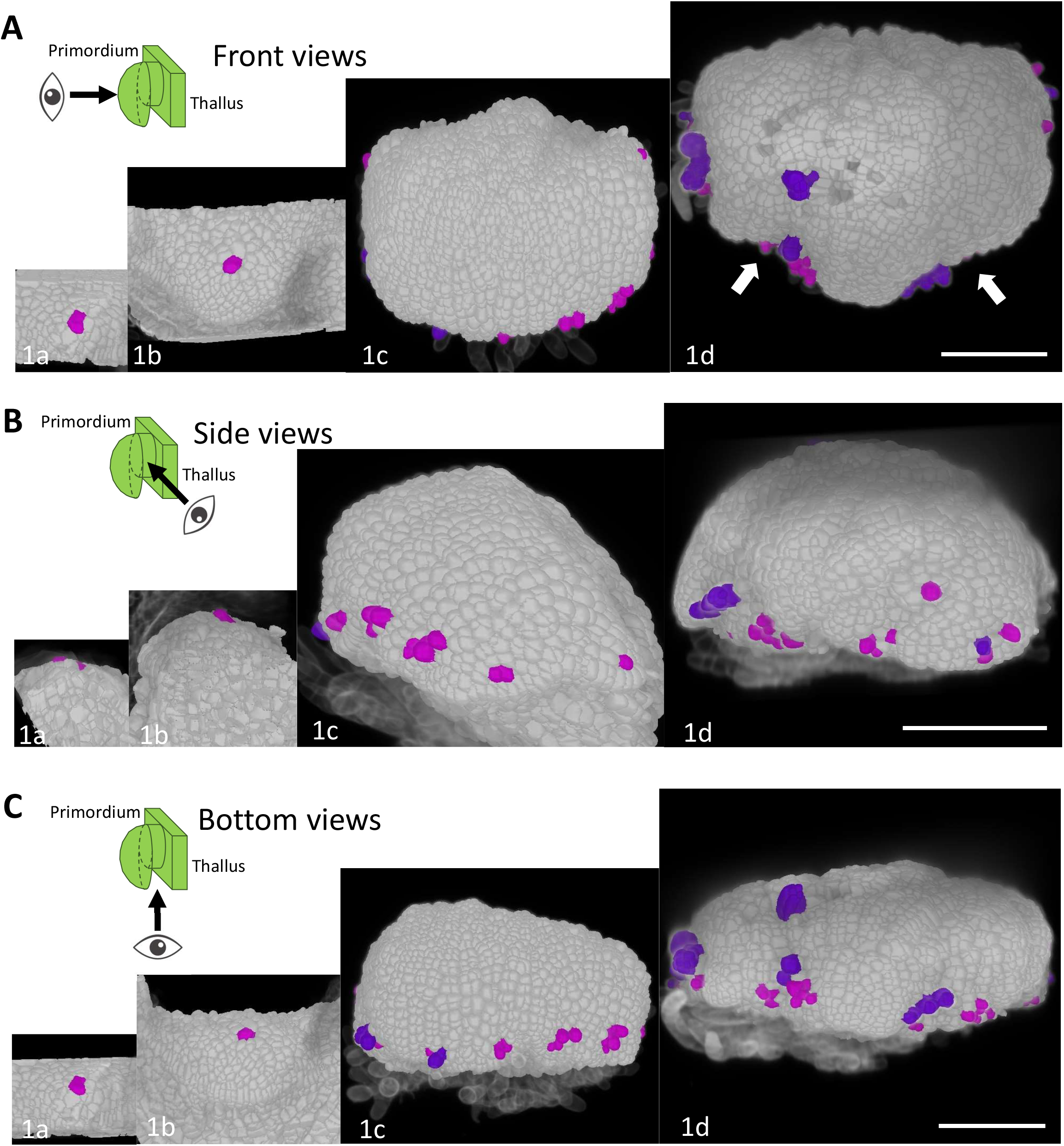
Morphogenesis and germline positioning during the archegoniophore development. Front views (A), side views (B) and bottom views (C) are shown for the primordia of different stages (1a-1e). Germline cells of early and late stages are colored pink and purple, respectively, according to Fig. 2E. White arrows in (A) indicate indentations. Primordia of stages 1a, 1b, 1c, and 1d were taken 3, 4, 6, and 7 days after the onset of FR irradiation, respectively. Scale bar, 100 µm.

**Figure 6.**
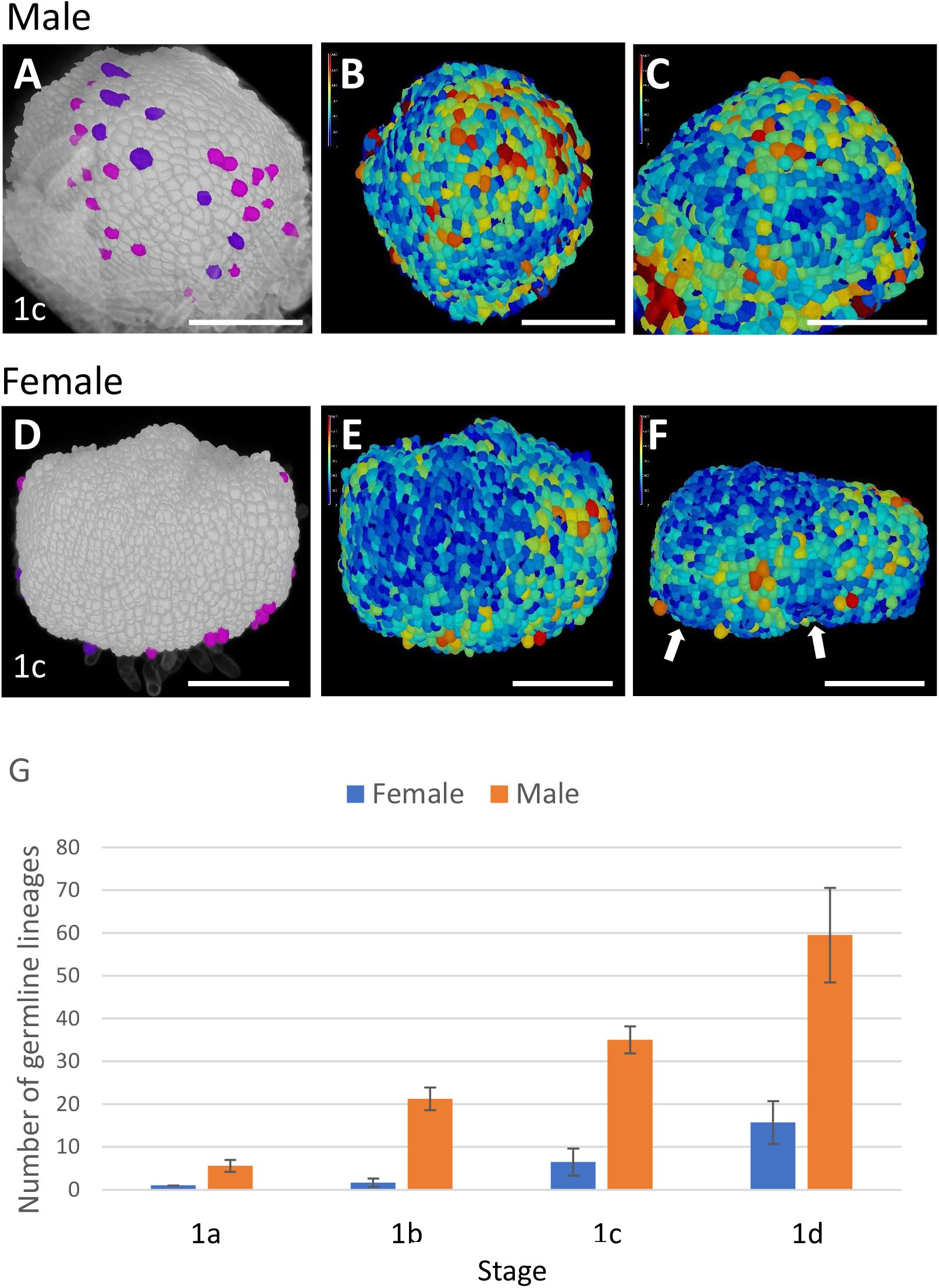
Cell volume distribution patterns and germline cell numbers of gametangiophore primordia. **(A-F)** 3D morphologies of antheridiophore (A-C) and archegoniophore (D-F) primordia and their cell size distribution patterns (A, B, D, E). Front views (A, B, D, E) and bottom views (C, F) are shown. White arrows in (F) indicate indentations. In (A) and (D), germline cells are colored pink and purple, respectively, according to Fig. 2E. **(G)** Number of germline cells in the primordia of different stages. *n*=3, 3, 13, 13 for the stage 1a, 1b, 1c, and 1d female primordia, respectively, and 7, 8, 5, 4 for the stage 1a, 1b, 1c, and 1d male primordia, respectively Scale bar, 100 µm.

**Figure 7.**
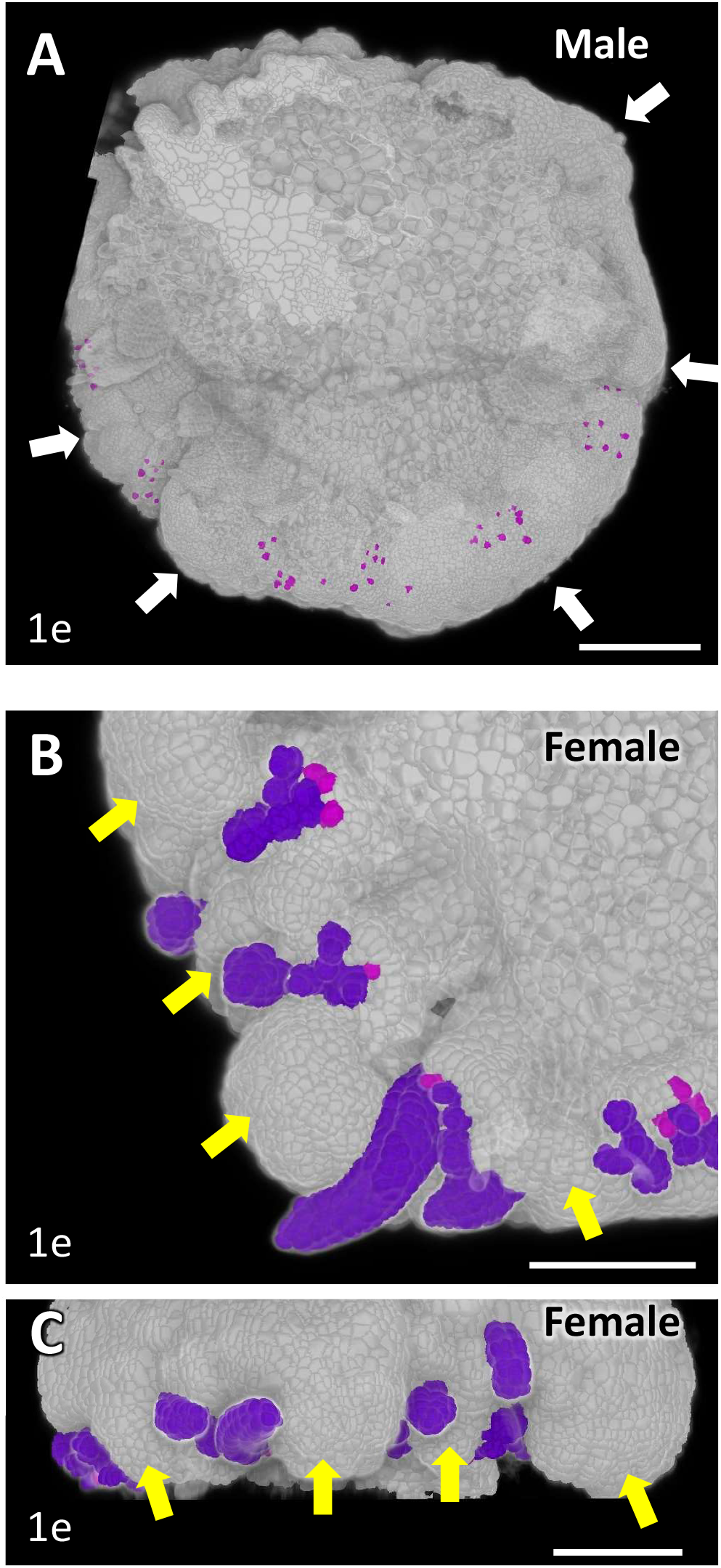
Development of sex-specific gametangiophore morphologies. Bottom views (A, B) and a side view (C) of 3D-reconstructed antheridiophore (A) and archegoniophore (B, C) primordia in the stage where sex-specific receptacle morphologies start to develop. White arrows in (A) and yellow arrows in (B) and (C) indicate incipient lobes and incipient finger-like rays, respectively. Germline cells of early and late stages are colored pink and purple, respectively, according to Fig. 2E. Scale bar, 100 µm.

In summary, early development of male and female gametangiophore primordia predicts the mature receptacle morphologies, though the primordia of these stages are yet ∼10 times smaller than mature gametangiophores. Beside the visible morphological differences, male and female primordia exhibit striking difference in the number and distribution patterns of germline lineages.

### Distinct cell proliferation and elongation may underlie the sex-specific gametangiophore morphogenesis

To elucidate the cell patterning processes that differentiate the male and female sexual morphologies in gametangiophore development, we analyzed the cell volume distribution using the 3D cell segmentation data of stage 1c gametangiophore primordia, where the distinct sexual morphologies became first recognizable (Figure 6A-6F). In male primordia, domains filled with small cells were found on the bottom surface and around the edge region (blue cells in Figure 6B and 6C; Supplementary video 5), whereas in the female primordium, small cells were preferentially distributed on the top surface and at the indentations (Figure 6E and 6F; Supplementary video 6). These observations suggest the existence of spatially distinct regulation of cell proliferation and/or elongation between male and female primordia at this stage.

### Mp*FGMYB* regulates female sexual differentiation from initial stage of gametangiophore development

Loss-of-function Mp*fgmyb* mutant females are masculinization in most sexual differentiation steps including gametangiophore morphogenesis, gametangium formation, and gamete differentiation (Hisanaga et al. 2019a). To investigate when and where Mp*FGMYB* functions during female sexual differentiation, we first carried out detailed 3D expression analyses of the transcriptional Mp*FGMYB* reporter in a wild-type female background, as well as the MpBNB-Citrine marker in the Mp*fgmyb* mutant background (Supplementary Figure S1). Consistent with the previous report (Hisanaga et al. 2019a), expression of Mp*FGMYB* was undetectable in the apical notch region of vegetative thalli (Figure 8A). After induction of reproductive growth by FR irradiation, the Mp*FGMYB* reporter started to express in archegoniophore primordia (Figure 8B; Supplementary video 9). Presumptive germline precursors that are destined to archegonia and protruded from the primordium surface exhibited stronger expression of the Mp*FGMYB* reporter in the following stage (Figure 8C; Supplementary video 10).

**Figure 8.**
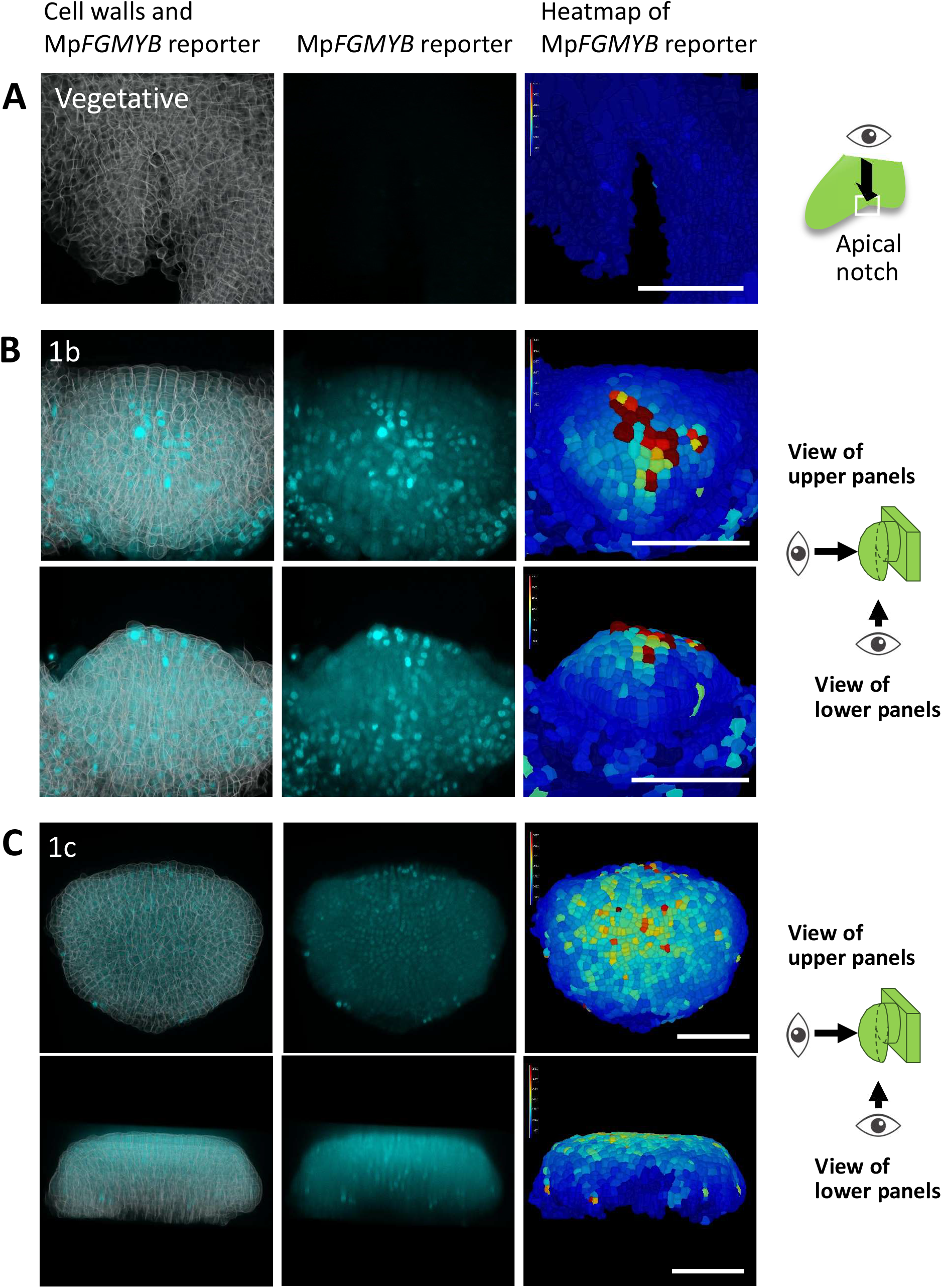
Expression pattern of Mp*FGMYB* transcriptional reporter as visualized in the reconstructed 3D morphology of archegoniophore primordia. **(A)** Top views of the apical notch region of a vegetative thallus, indicating the absence of Mp*FGMYB* expression. **(B, C)** Expression patterns of *MpFGMYB* reporter in archegoniophore primordia of different stages. Front views and bottom views are shown for the identical samples for each stage. Scale bar, 100 µm

Gametangiophore morphogenesis and germline precursor positioning of Mp*fgmyb* mutants closely followed those of wild-type males. Multiple MpBNB-expressing gametangium initials were found on the slightly convex surface of incipient gametangiophore primordia emerged from the apical notch region after FR irradiation (Figure 9, stage 1a; Supplementary video 11). In later stages, gametangiophore primordia expanded to have a dome-like morphology and the number of germline precursors increased (Figure 9, stage 1b). As the receptacle primordia grew larger, they gradually became flattened as in wild-type males. Developing gametangia (Figure 9, purple) were observed on the upper surface of receptacle primordia with more mature gametangia located closer to the center of the receptacle primordia as in wild-type males (compare Figure 4 and 9, stage 1d). Thus, Mp*fgmyb* is masculinized from the earliest observable stage of sexual dimorphism development, supporting the notion that Mp*FGMYB* is a master regulator of female sexual differentiation in *M. polymorpha*.

**Figure 9.**
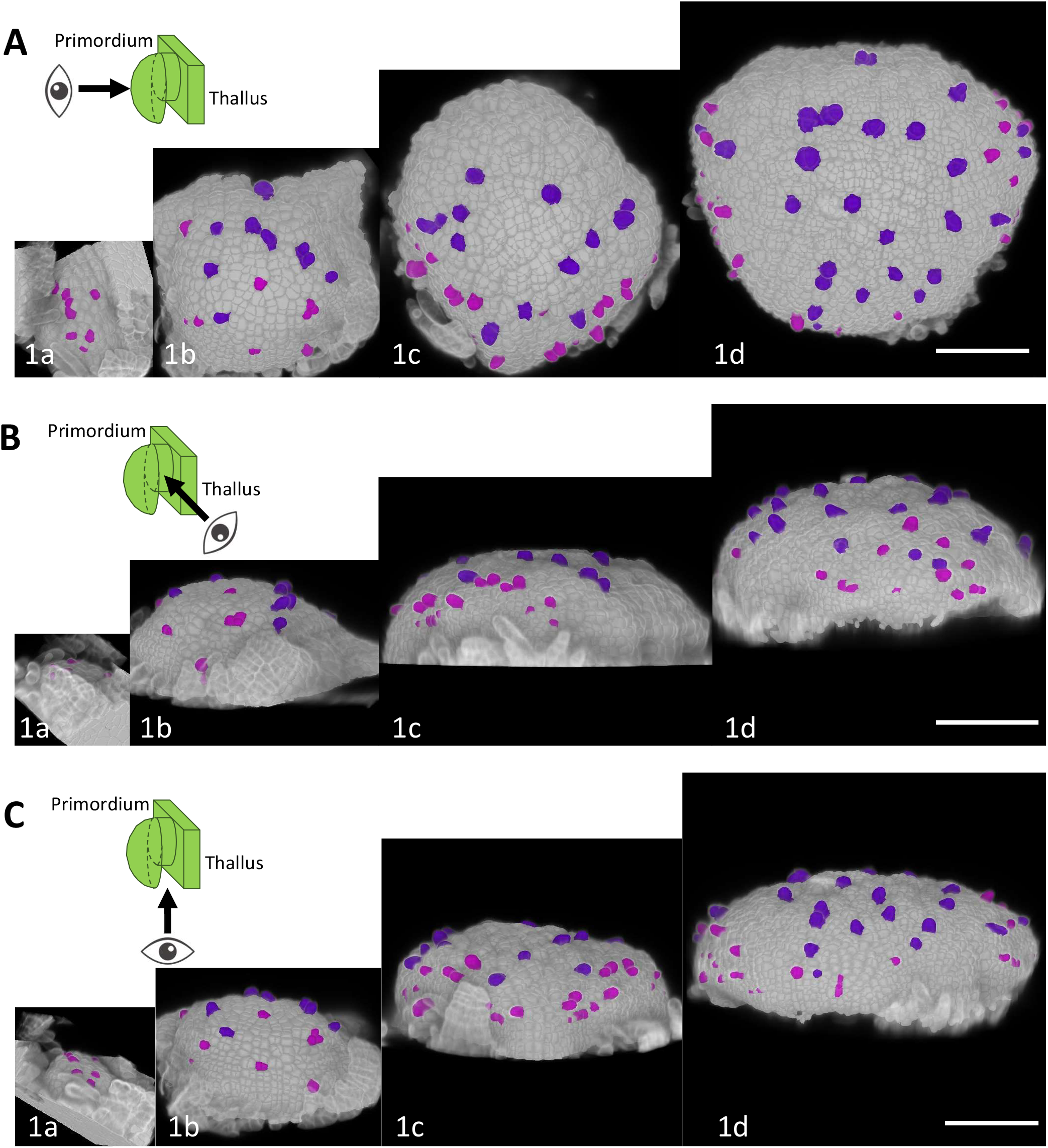
Morphogenesis and germline positioning during the development of masculinized gametangiophore primordia of Mp*fgmyb*. Front views (A), side views (B), and bottom views (C) are shown for the primordia of different stages (1a-1d). Germline cells of early and late stages are colored pink and purple, respectively, according to Fig. 2E. Primordia of stages 1a, 1b, 1c and 1d were taken 2, 3, 4 and 5 days after the onset of FR irradiation, respectively. Scale bar, 100 µm.

## Discussion

### Growth phase transition and germline differentiation are coupled in *M. polymorpha*

Sexual dimorphism of mature gametangiophores of *M. polymorpha* has been documented since the mid-15th century (reviewed by Bowman 2016), yet how its characteristic sexual branch dimorphism is established has not been explicitly described at cellular resolution. In this study, we performed cell-level 3D-morphological analysis using the MorphoGraphX (Strauss et al. 2022; Vijayan et al. 2021) to address this question. Our data revealed strong correlation between spatial distribution patterns of germline precursors and morphogenic processes of gametangiophore receptacles. In both males and females, MpBNB-expressing germline precursors first emerged as early as in the stage where incipient gametangiophore primordium could be barely recognized as a slight protrusion in the apical notch region. More germline precursors repeatedly emerged as the gametangiophore primordia expand, but with different frequencies and spatial patterns between males and females. These observations indicate that sexual morphogenesis of gametangiophore and specification of germline precursors occur in parallel. This is in good contrast with the reproductive development of sporophyte-dominant flowering plants, where conspicuous sexual morphogenesis first takes place in sporophytic floral organs (stamens and carpels), followed by meiocytes differentiation (Barrett and Hough 2013). Previous studies indicated that specification of male and female gametes requires sporophyte-derived factors in flowering plants (Olmedo-Monfil et al. 2010; Tidy et al. 2022; Zhao et al. 2001; Zhao et al. 2017). Thus, sporophytic sexual differentiation is a prerequisite for precise specification of germline precursors. By contrast, in gametophyte-dominant *M. polymorpha*, differentiation of sexual morphologies and germline specification appear to be spatiotemporally coupled to each other.

Then, how is the progression of germline specification and sexual branch formation coordinately regulated? Our data suggest that Mp*BNB* plays an important role. Loss-of-function Mp*bnb* mutants are unable to initiate sexual branch formation (Yamaoka et al. 2018), though MpBNB-Citrine reporter was found to express transiently in germline precursors, not in entire primordia (Yamaoka et al. 2018). Our detailed 3D visualization confirmed the germline precursor-specific expression of MpBNB throughout the course of early gametangiophore development, suggesting that Mp*BNB* primarily acts in germline differentiation, which in turn promotes sexual branch formation by unknown mechanisms. In this scenario, sexual morphogenesis follows germline differentiation in *M. polymorpha*, an order opposite to that in flowering plants, but analogous to early germline segregation and sexual organ differentiation in animal development (Lanfear 2018).

### Different germline positioning may underlie sex-specific gametangiophore morphologies

Our 3D morphological analysis revealed that male and female gametangiophore primordia initially exhibit a similar dome-like morphology. At this stage, however, spatial distribution patterns of Mp*BNB*-expressing germline initials (which later develop into gametangia) are already different between males and females. This difference is even apparent in the initial stage where gametangiophore primordia can be barely recognized as a slight protrusion in the apical notch region. In males, many germline precursors emerge on the convex surface of incipient primordia and their number increases as the primordia expand, whereas in females significantly fewer germline precursors are initially formed, and their sparse distribution pattern is maintained in later stages. Importantly, spatial distribution patterns of germline precursors correlated well with the sexual morphologies of male and female gametangiophores. In males, germline precursors are scattered over the uppers surface of antheridiophore primordia and develop into antheridia on the top surface of disc-shaped antheridiophore receptacles. In females, germline precursors are localized along the peripheral region of archegoniophore primordia with regular intervals, and eventually develop into mature archegonia. Cells occupying the space between the two neighboring germline precursor clusters proliferate to form protrusions, which later develops into the finger-like rays seen in mature archegoniophores. Thus, germline precursor positioning appears to play an important role in establishing the sexual dimorphism of gametangiophore receptacles of *M. polymorpha* (Figure 10).

**Figure 10.**
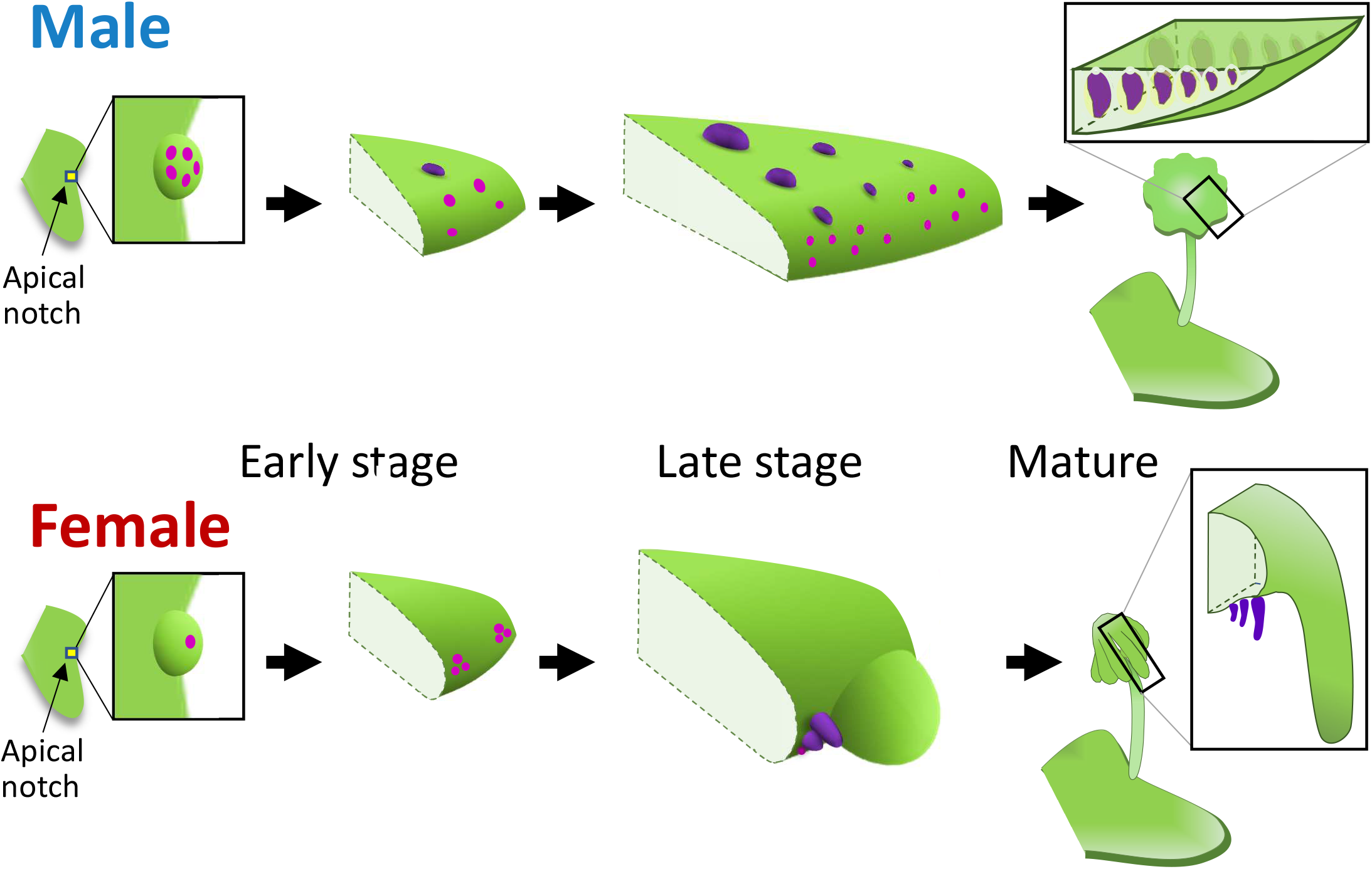
Diagrams illustrating the process of sexual dimorphism development in *M. polymorpha*. After induction of reproductive growth, germline precursors (pink) differentiate on the slightly convex surface of incipient receptacle primordia in the apical notch region. In this stage little morphological difference is apparent between male and female receptacles, with both having a dome-like shape, whereas they show different spatial arrangement in the germline precursors. As development proceeds, male receptacles (top) become flattened and the antheridium primordia (purple) develop over the top surface of receptacles, which later acquire the characteristic antheridiophore morphology with peripheral lobes. In females (bottom), archegoniophore receptacles (bottom) develop archegonium primordia (purple) along the receptacle periphery with regular spacing. Later, regions between adjacent archegonia clusters extend to develop finger-like rays to confer the characteristic receptacle morphology. Female plants lacking Mp*FGMYB* precisely follow the male-type developmental processes both in germline precursor placement and receptacle morphogenesis, indicating a pivotal role of MpF*GMYB* in female sexual differentiation.

### Mp*FGMYB* promotes female sexual differentiation from the initial stage of reproductive development

Mp*FGMYB* encoding a Myb-type transcription factor has been identified as a key regulator of female sexual differentiation in *M. polymorpha* (Hisanaga et al. 2019a). When Mp*FGMYB* is knocked out in females, their sexual morphologies were masculinized in multiple scales, including gametangiophore morphologies, gametangium differentiation, and gamete formation. This previous study indicated that the default sexual differentiation program of *M. polymorpha* is for males, and that Mp*FGMYB* “overwrites” it with a female sexual differentiation program. While the previous study revealed the expression of the Mp*FGMYB* reporter in the apical notch region of thalli after FR-irradiation, it has been unknown from what stage of reproductive development Mp*FGMYB* functions to promote female sexual morphogenesis.

Our 3D expression and phenotypic analyses clearly indicated that Mp*FGMYB* is required from the early stage of sexual dimorphism development. In genetically female Mp*fgmyb* mutants, the MpBNB-expressing germline precursors were scattered over early gametangiophore primordia as seen in wild-type males. The Mp*FGMYB* reporter was expressed in several layers over the entire surface of archegoniophore primordia in wild-type females, with a few protruding cells destined to archegonia exhibiting significantly higher expression. These observations suggest that Mp*FGMYB* functions to regulate germline positioning at the initial stage of archegoniophore development, in addition to gametangia morphogenesis that takes place in more later stages. These results are consistent with the previously proposed role of MpFGMYB as a master regulator of female sexual differentiation in *M. polymorpha* (Hisanaga et al. 2019a).

## Materials and Methods

### Plant materials and growth condition

Wild-type female and male strains of *M. polymorpha* used in this study were Takaragaike-2 (Tak-2) and Takaragaike-1 (Tak-1), respectively (Ishizaki et al. 2016). Mp*fgmyb* knock-out plants harboring the MpBNB-Citrine marker was generated by transforming the female Mp*BNB-Citrine* knock-in plants (Yamaoka et al. 2018) with the previously described gRNA construct *pMpGE011_MpFGMYBge01* (Hisanaga et al. 2019a). The transcriptional Mp*FGMYB* reporter line was prepared by transforming the *MpFGMYBpro:H2B-mNeonGreen:MpSUF* construct describe below to the wild-type Tak-2 plants as described previously (Tsuboyama et al. 2018).

Plants were cultured on a half-strength Gamborg’s B5 agar medium under continuous white light at 22 °C and asexually propagated through gemmae. To induce reproductive growth, gemmae pre-cultured for 10 days were transferred to vermiculite-containing pots and illuminated with continuous white light supplemented with FR as described previously (Hisanaga et al. 2019a).

### DNA construction

*MpFGMYBpro:H2B-mNeonGreen:MpSUF* was constructed as follows using the primers listed in Supplementary Table 1. Mp*FGMYB* and Mp*SUF* sequences were amplified from Tak-1 genomic DNA using primer pairs of pENTR1a-AscI-MpFGMYB4kbup-F and pENTR1a-MpFGMYBPmeI-R, and MpFGMYBPmeI-F and pENTR1a-MpSUF5kbup-R, respectively. The Mp*FGMYB* sequence was subcloned into an EcoRI site of pENTR1A vector (Thermo Fisher Scientific, MA) using the In-Fusion system (Takara Bio, Shiga, Japan). The resultant Mp*FGMYB* plasmid was digested with PmeI and NotI, and the amplified Mp*SUF* sequence was ligated into the PmeI/NotI sites of Mp*FGMYB* plasmid using the In-Fusion system to create *pENTR1a-MpFGMYB-MpSUF* carrying the entire sequence of the Mp*FGMYB*/Mp*SUF* gene locus including 4-kb upstream and 5.3-kb downstream regions. Next, to replace the Mp*FGMYB* coding sequence with a reporter sequence *H2B-mNeonGreen*, two partial sequences of *pENTR1a-MpFGMYB-MpSUF* were amplified using primer pairs of MpFGMYB4kbup-Cloning-F and PmeI-MpFGMYBpro-Rv, and PmeI-MpFGMYBpro-Fw and MpFGMYBEco105I-R, and combined into a single sequence by overlap extension PCR, which was then inserted into the AscI/Eco105 sites of pENTR1a-MpFGMYB-MpSUF to create pENTR1a-MpFGMYBpro-PmeI-MpSUF. The coding sequence of *AtHTB1* (AT1G07790) was amplified from *pKI-GWB2 H2B* vector (provided by Dr. Kimitsune Ishizaki) using a primer pair of AtHTB1-Cloning-F-dTOPO and AtHTB1-GGSGGS-R. The coding sequence of *Arabidopsis*-codon-optimized mNeonGreen was amplified from pENTR Atco_mNeonGreen vector (provided by Dr. Ryuichi Nishihama) using a primer pair of GGSGGS-Atco_mNG-F and Atco_mNG-Cloning-R. These sequences were combined into a single sequence by overlap extension PCR, and subcloned into pENTR/D-TOPO vector (Thermo Fisher Scientific). The resultant vector was used to amplify the coding sequence of *H2B-mNeonGreen* using a primer pair of MpFGMYBp-AtHTB1-Fw and MpFGMYBt-mNG-Rv, and the resultant sequence was ligated into the PmeI site of pENTR1a-MpFGMYBpro-PmeI-MpSUF using a SLiCE reaction (Zhang et al. 2012). The resultant entry vector was used for a recombination reaction with pMpGWB101 (Ishizaki et al. 2015) using Gateway LR clonase II Enzyme mix (Thermo Fisher Scientific) to create *pMpGWB101-MpFGMYBpro:mNeonGreen:MpSUF*.

### Histology and microscopy

Gametangiophore primordia were dissected using forceps and needles under a dissecting microscope and fixed with a 1% (w/v) formaldehyde (FA) solution for 1 hour followed by washing three times with PBS buffer. Samples were cleared in a decolorization solution [100 mM sodium phosphate buffer pH 8.0, 20% (w/v) caprylyl sulfobetaine, 7.5% (w/v) sodium deoxycholate] overnight. After washing in PBS, samples were stained with 0.1% (v/v) Renaissance 2200 (Renaissance Chemicals, Selby, UK; Musielak, 2015) in PBS for 10 min. After washing in PBS, samples were incubated in a series of mounting solutions [20%, 50% and 70% (w/w) iohexol in PBS] (Kurihara et al. 2015; Sakamoto et al. 2022), and observed using a Leica TCS SP8 confocal laser scanning microscope (Leica Microsystems, Wetzlar, Germany). Typically, Z-stack images were collected in 0.2-0.3 µm increments for the depth of 70-180 µm to construct the 3D images.

### Cell segmentation analysis using MorphoGraphX

Cell segmentation was performed using the MorphoGraphX (MGX) program as described previously (de Reuille et al., 2015, Vijayan, 2021). Briefly, confocal stack images of Renaissance 2200-stained gametangiophore primordia were first enhanced for the image contrast using the Stack Contrast Adjustment Plugin of the ImageJ software (Čapek et al. 2006). After loading to MGX, images were blurred by a Gaussian filter with a radius of 0.8 µm in xyz. Segmentation was carried out by the ITK morphological watershed function with the default threshold of 3000, and 3D cell meshes were generated from the segmented image stacks using the Marching Cubes 3D function with a 2.5 µm cube size. For germline precursor labeling, the cell meshes corresponding to the MpBNB-Citrine expressing cells were labeled using the parent label function of MGX.

## Acknowledgement

We thank Masako Kanda for technical assistance, and Kimitsune Ishizaki and Ryuichi Nishihama for materials. YC was supported by the MEXT scholarship.

---

**Supplementary Figure 1.**
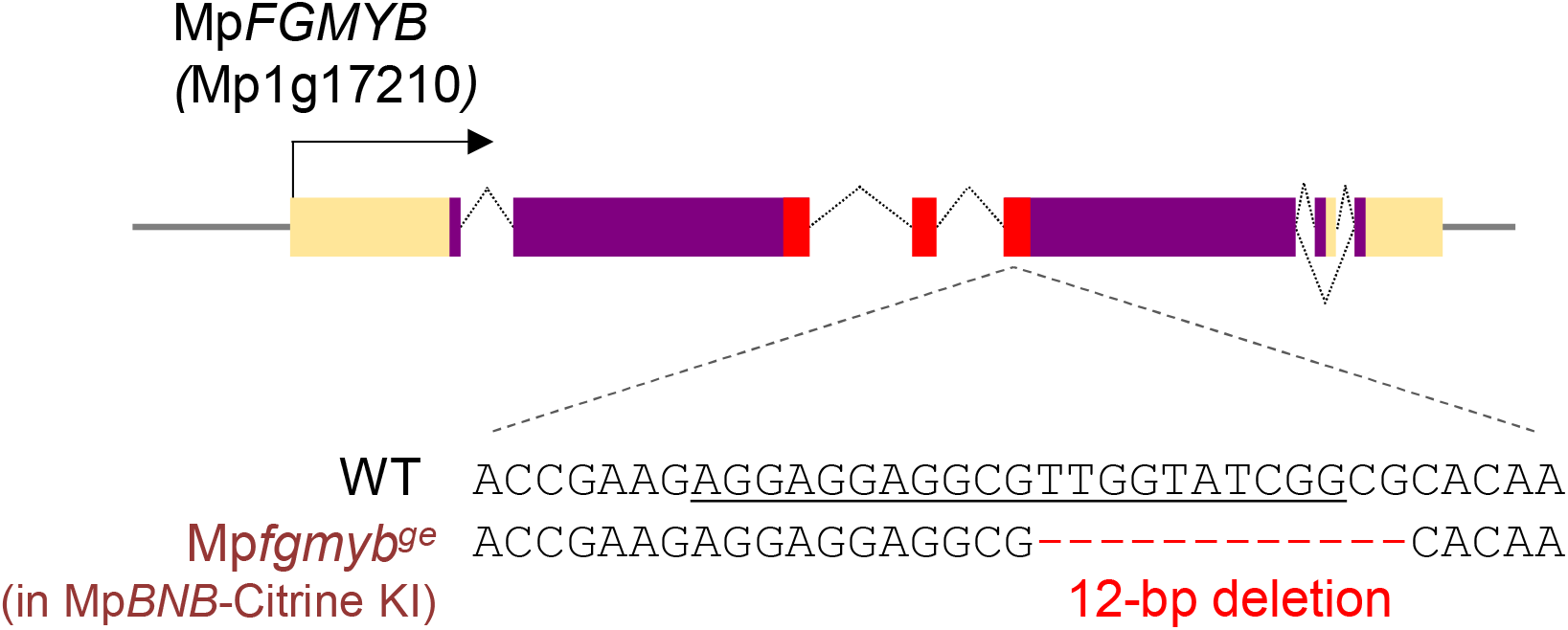
CRIPSR-induced Mp*fgmyb* mutant in the Mp*BNB*-Citrine knock-in line. Orange, purple, and red boxes in the gene diagram indicate untranslated regions, protein-coding regions, and Myb domain-coding parts, respectively. Alternative splicing is represented by folded lines connecting the exons. Underline indicates the gRNA target sequence.

